# Synonymous and non-synonymous transitions/transversions vividly disclose purifying selection in *Escherichia coli* coding sequences

**DOI:** 10.1101/2022.11.03.515082

**Authors:** Pratyush Kumar Beura, Ruksana Aziz, Piyali Sen, Saurav Das, Nima Dondu Namsa, Edward J Feil, Siddharatha Sankar Satapathy, Suvendra Kumar Ray

**Affiliations:** Department of Molecular Biology and Biotechnology, Tezpur University, Tezpur, Assam, India, 784028; Department of Computer Science and Engineering, Tezpur University, Tezpur, Assam, India, 784028; Centre for Multidisciplinary Research, Tezpur University, Tezpur, Assam, India, 784028; The Milner Centre for Evolution, Department of Biology and Biochemistry, University of Bath, Bath BA2 7AY, United Kingdom

**Keywords:** substitution mutation, single nucleotide variations, transition to transversion ratio, codon degeneracy, pretermination codon, purifying selection

## Abstract

Transition (*ti*) and transversion (*tv*) are the major causes for genome variation. The accurate estimation of *ti* to *tv* ratio 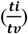 in genomes is crucial for understanding of mutational and selection processes in organisms as it is influenced by both codon degeneracy and pretermination codons (PTC). Therefore, we developed a method (accessible at https://github.com/CBBILAB/CBBI.git) to estimate 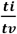 ratio by accounting codon degeneracy as well as PTC in protein coding sequences. Our findings revealed a distinct impact of codon degeneracy and PTC on the 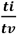 ratio in the *Escherichia coli* genome. We observed a decreasing order among the frequencies of different base substitutions such as synonymous transition (*Sti*) > synonymous transversion (*Stv*) > non-synonymous transition (*Nti*) > non-synonymous transversion (*Ntv*) in *E. coli* genome. The correlation was strong between *Sti* and *Stv* values (Pearson *r* value 0.795) whereas the correlation was weak between *Sti* and *Nti* (Pearson *r* value 0.192). Coding sequences with similar *Sti* values exhibited a wide range of *Nti* values. This indicated the varying strength of purifying selection acting on the coding sequences. In concordance with the assumption, the genes having higher *Nti* values were observed with lower codon adaptation index (CAI) values than that of the genes having lower *Nti* values. Our approach is convenient to visualize the frequency of base substitution variation as well as selection in protein coding sequences. The proposed method is useful to estimate different 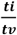 ratios accurately in coding sequences and is insightful from an evolutionary perspective.

**Article Summary:** Genetic diversity is pivotal in evolution, with base substitution as a key driver. Transition (*ti*) frequency surpasses transversion (*tv*) frequency in genomes, making 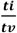 ratios a valuable metric for studying mutation bias. Our improved estimator for 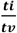 calculation accounts for codon degeneracy and nonsense substitutions in pretermination codons. Additionally, we unveil insights into the frequency of different substitutions such as *Sti, Stv, Nti*, and *Ntv* and demonstrate the impact of selection on protein coding sequences.

## Introduction

The degeneracy in the genetic code table is an important feature to study molecular evolution (Crick *et al*. 1961; Khorana *et al*. 1966). The different degenerate codons influence the transition (*ti*) to transversion (*tv*) ratio 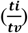 as follows: synonymous variation in a *two-fold degenerate* (TFD) codon occurs *via* only one transition whereas the same in a *four-fold degenerate* (FFD) codon occurs via one transition and two transversions (Supplementary Fig. 1a). In addition to the above, compositional variation of pretermination codons (PTC) (Supplementary Fig. 1b and 1c) (Modiano *et al*. 1981) among the coding sequences is also likely to influence the 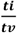 ratio because non-synonymous variations leading to termination codons purged out rapidly in a population due to stronger purifying selection (Li *et al*. 1981; Eynden *et al*. 2016; Morales *et al*. 2021). Considering in general, a *ti* being more frequent than a *tv* (Seplyarskiy *et al*. 2012; Duchêne *et al*. 2015; Stoltzfus and Norris 2016; Lewis *et al*. 2016; Lyons and Lauring 2017; Schroeder *et al*. 2017; Sen *et al*. 2022) which is known as Kimura’s two parameter model (Kimura 1980), and a stronger purifying selection on non-synonymous variation than a synonymous variation (Hurst and Pál 2001), the 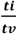 ratios of TFD codons and FFD codons are expected to be different. Accordingly, it has been recently reported that the synonymous variation in TFD codons is relatively higher than the FFD codons in *Escherichia coli* (Aziz *et al*. 2022). Therefore, in any organism, the 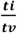 ratio is likely to vary across the coding sequences having differences in degenerate codon composition (Beura *et al*. 2023). Hence, an estimation of 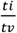 ratio in a coding sequence by accounting codon degeneracy and PTC compositions will be important to further carrying out comparative analysis of this value among coding sequences in a genome.

**Fig. 1.**
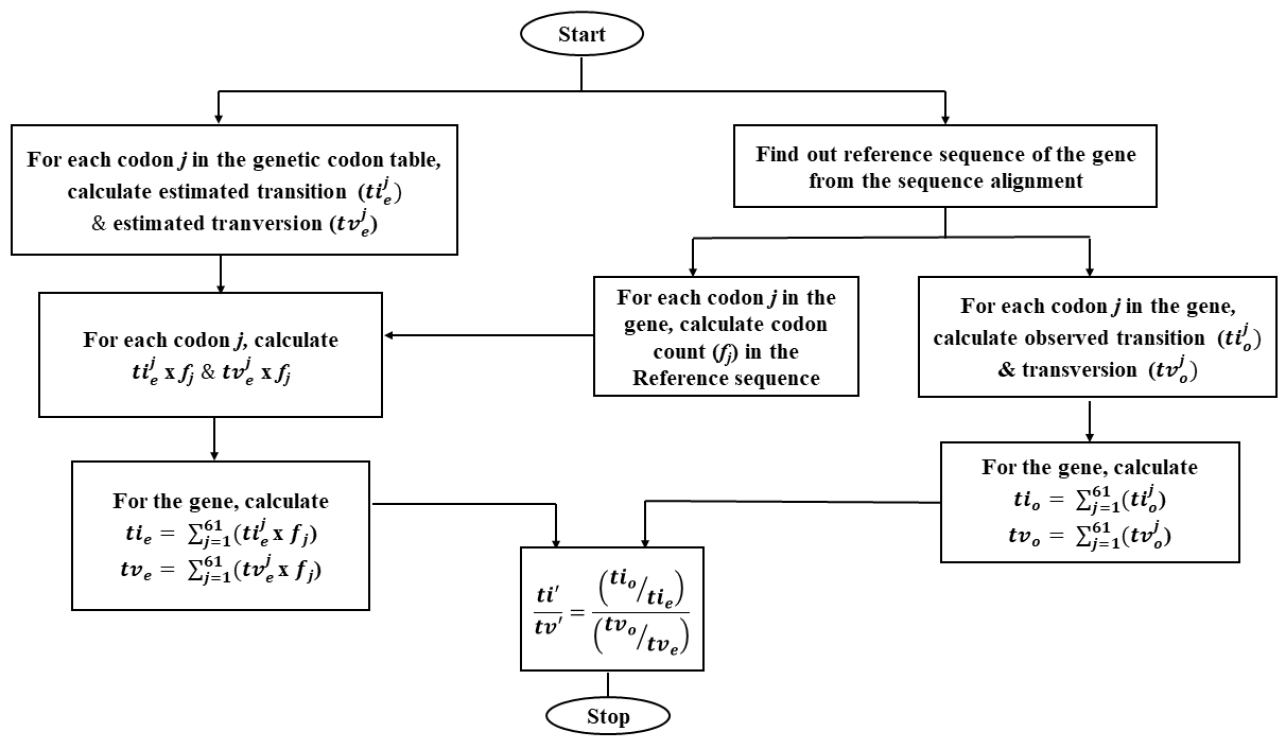
Schematic representation demonstrating step wise workflow for calculation of 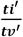 using the improved method. Step wise calculations for the workflow are fgiven for a sample gene in the Supplementary Table S5, S6, S7.

Considering the number of possibilities of the two types of base variations in genomic DNA (Fersht and Knill-Jones 1981) (Supplementary Fig. 1d). The ratio of *ti* to *tv* is theoretically expected to be 0.5. But the ratio is generally observed more than 0.5 in the genomes (Duchêne *et al*. 2015; Stoltzfus and Norris 2016). Because of the following factors such as geometry of DNA double helix (Topal and Fresco 1976), modification of DNA bases such as deamination of cytosine and adenine (Kino and Sugiyama 2001; Rocha *et al*. 2006; Bhagwat *et al*. 2016), secondary structure in the RNA (Sen *et al*. 2022) and codon degeneracy are known to elevate *ti* phenomenon in a genome over *tv* (Muse and Gaut 1994; Aziz *et al*. 2022). In a standard genetic code table out of 549 total single nucleotide variations (SNVs) involving 61 sense codons (9 SNVs per codon), only 134 SNVs are synonymous (24.4%). Among synonymous (*Sti* and *Stv*) and non-synonymous (*Nti* and *Ntv*) variations, the theoretically estimated (_e_) numbers of *Sti, Stv, Nti* and *Ntv* in a genetic code table are 62, 72, 121 and 294, respectively (Supplementary Table 1). Yet the observed synonymous variations are always higher than the expected proportion across genes (Wolfe *et al*. 1989; Tamura 1992; Moriyama and Powell 1997; Ngandu *et al*. 2008) due to stronger purifying selection on non-synonymous variations than the synonymous variations (Hurst and Pál 2001). Hence, by accounting the codon degeneracy as well as PTC, the *Sti, Stv, Nti* and *Ntv* in the genetic code table can be estimated theoretically for each codon (Supplementary Table 2).

**Table 1.**
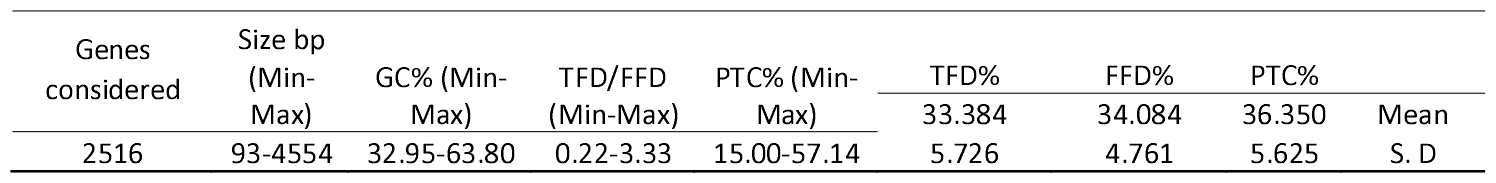
Nucleotide and codon compositional features of the 2516 coding sequences considered in the study.

**Table 2.**
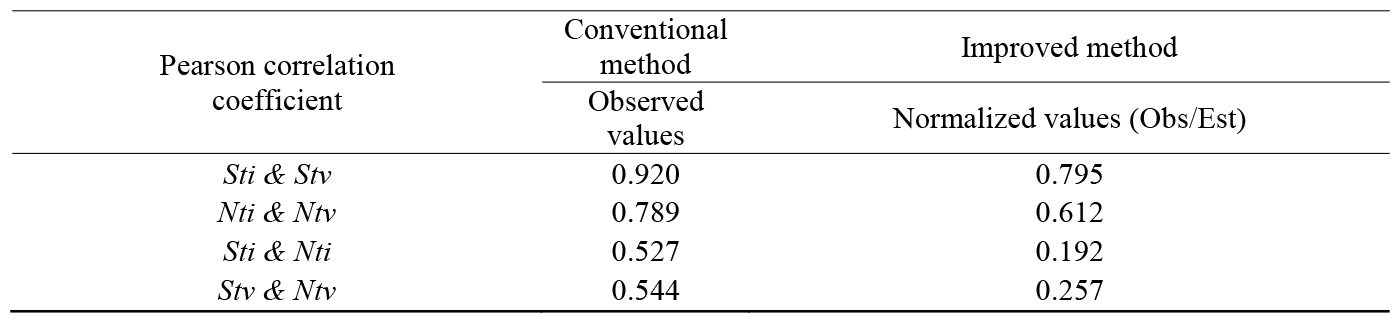
Pairwise Pearson correlation coefficient among *Sti, Stv, Nti* and *Ntv* through conventional and improved estimator.

| Pearson correlation<br>coefficient | Conventional<br>method | Improved method |
| --- | --- | --- |
|  | Observed<br>values | Normalized values (Obs/Est) |
| *Sti* & *Stv* | 0.920 | 0.795 |
| *Nti* & *Ntv* | 0.789 | 0.612 |
| *Sti* & *Nti* | 0.527 | 0.192 |
| *Stv* & *Ntv* | 0.544 | 0.257 |

The codon substitution model suggests that sites within codons evolve at different rates and consequently they should not be equally treated (Muse and Gaut 1994; Shapiro *et al*. 2006; Arenas 2015). In the codon substitution model, using likelihood approach the 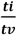 ratio was estimated in accordance with the codon degeneracy class which was implemented in estimating *dN*/*dS* in coding sequences (Muse and Gaut 1994; Goldman and Yang 1994). Recently, by accounting 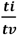 substitution rate difference in codons as well as nonsense variations in PTC, the *dN*/*dS* values have been improved (Aziz *et al*. 2022). This finding implies that factors such as codon degeneracy and PTC composition should be considered while analyzing the evolution of protein-coding sequences. The ratio of *ti* to *tv* has been studied for reliable estimation of sequence distance and phylogeny reconstruction (Yang and Yoder 1994). Using maximum likelihood approach, the 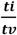 ratio (commonly known as kappa κ) has been used to mainly compare the 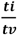 values among different species (Yang and Yoder 1994). The parameter κ has not been implemented to compare coding sequences within a genome in the context of their compositional differences of different degenerate classes before. As the selection pressure on synonymous and non-synonymous variations are heterogeneous (Bartolomé *et al*. 2005), it is important to analyze the 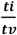values among different coding sequences within an organism by accounting synonymous sites and non-synonymous sites regarding their compositional differences of different degenerate codons.

It is noteworthy that in a genome, the number of potential replacements is greater for non-synonymous alterations, whereas the frequency of actual variations is higher for synonymous alterations among coding sequences. The proposed improved estimators account for the observed variations out of the total possible variations. On the contrary, the conventional method only takes an account of the observed *ti* and *tv*. The possible sites for *Sti* and *Stv* are also different among coding sequences, which is dependent upon the codon composition belonging to different degeneracy classes. The conventional 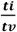 ratio does not adequately account for the limitations of variable selection against synonymous and non-synonymous changes. Given that there is a significant variation of substitution rates among protein coding sequences within a genome (Wu and Li 1985), an appropriate methodology for the estimation of *ti* to *tv* ratio is required. We are presenting an improved estimator that builds upon the research conducted by Yasuo Ina in 1998 (Ina 1998). Considering the differential selective constraint in synonymous and non-synonymous variations it is crucial to separately analyze the synonymous 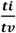 and non-synonymous 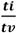 to gain a comprehensive understanding of the overall 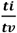 Hence, all these parameters make the requirement of an improved estimator instead of the conventional estimator. The proposed estimators used in this study are helpful to estimate the *ti* and *tv* rate differences in coding sequences by accounting the compositional difference in terms of codon degeneracy and PTC. The proposed estimator is useful in comparative study of mutation bias between two coding sequences in an organism, as it normalizes the 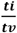 ratio in form of observed variations out of total possible variations. Further, we observed a correlation between TFD: FFD and 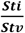 values and the PTC% and 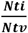 values. We tried to establish the biological significance of codon composition and 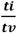 ratio across 2516 coding sequences of *E. coli*. Interestingly, coding sequences with similar synonymous variation frequency are found to have a wide variation regarding their non-synonymous variation frequency. The improved method developed in the present study may be applied to study different 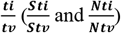 across coding sequences.

## Materials and Methods

### Coding sequences of *E. coli* genome and finding out *ti* and *tv*

In this study, we carried out a computational analysis of SNVs in 2516 protein coding sequences across 157 strains of *E. coli* (Thrope *et al*. 2017). The information regarding the dataset selected for the present study is provided in Supplementary Table 3. Although the public database has ∼4500 coding sequences in *E. coli* genome, we excluded those coding sequences exhibiting size differences and improper alignment/annotation across the strains. Codons having double or triple substitutions or variations represented by an ambiguous nucleotide were not considered further in the analyses of the variations. The procedure in detail used for estimating SNVs is represented in Supplementary Table 4 using example of a hypothetical sequence. This procedure is based on intra-species genome sequence comparison to find SNVs in bacterial genomes. A reference sequence was generated considering the most frequent nucleotide occurrence at a site in the gene sequence. The logic behind the derivation of the reference sequence is that the most abundant nucleotide in a position is considered as the ancestral nucleotide for the position (Sen et al. 2022; Aziz *et al*. 2022; Sen 2022, Beura *et al*. 2023).The observed SNVs (SNV_o_) at different positions of each codon were categorized into *Sti*_*o*_, *Stv*_*o*_, *Nti*_*o*_ and *Ntv*_*o*_, where S for synonymous, N for non-synonymous, *ti*_*o*_ for observed transition and *tv*_*o*_ for observed transversion, and subsequently *ti*_*o*_ (*Sti*_*o* +_ *Nti*_*o*_) and *tv*_*o*_ (*Stv*_*o*+_ *Ntv*_*o*_) for each gene sequence were calculated (Supplementary Table 5). After finding out the observed variation values for a gene, we estimated the total number of variations such as *Sti*_*e*_, *Stv*_*e*_ *Nti*_*e*_, *Ntv*_*e*_, *ti*_*e*_ and *tv*_*e*_ possible theoretically in the gene. We derived the reference sequence for each gene based on the methodology explained above. The reference sequence was used for the theoretical estimation of these variations. For example, if GGG codon abundance is 10 in a gene, then the total theoretically estimated value for *Sti, Stv, Nti* and *Ntv* are 10, 20, 20 and 40 (*Sti, Stv, Nti* and *Ntv* for GGG codon is 1, 2, 2, and 4, respectively). The *Sti, Stv, Nti*, and *Ntv* for each codon in the genetic code table are given in Supplementary Figs (1a, 1b, 1c). This information for each codon was used to calculate the theoretically estimated SNVs (*Sti*_*e*_, *Stv*_*e*_, *Nti*_*e*_ and *Ntv*_*e*_) for each gene under the study (Supplementary Table 6). The values of the observed SNVs (*Sti*_*o*_, *Stv*_*o*_, *Nti*_*o*_, *Ntv*_*o*_, *ti*_*o*_, and *tv*_*o*_) and the values of estimated SNVs (*Sti*_*e*_, *Stv*_*e*_ *Nti*_*e*_, *Ntv*_*e*_, *ti*_*e*_, and *tv*_*e*_) of a coding sequence are then used in the modified estimator to find out different 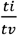 ratio 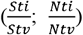 (Supplementary Table 7). A detailed workflow is provided in Fig. 1.

### Improved estimators

We proposed an improved mathematical equation for the calculation of different 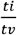 ratio as described below.

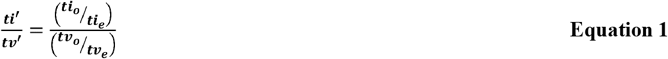

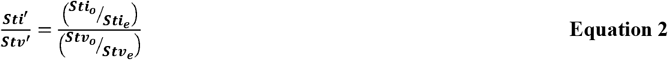

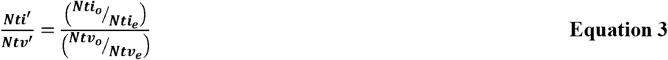

Where, 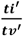 is improved 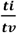 ratio, 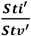 is improved 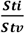 ratio, 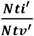 is improved 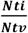 ratio: *ti*_*o*_ is number of transitions observed, ***ti***_***e***_ is the number of possible transitions estimated theoretically, *tv*_*o*_ is the number of transversions observed, *tv*_*e*_ is the number of possible transversions estimated theoretically, ***Sti***_***o***_ is number of synonymous transitions observed, ***Sti***_***e***_ is the number of theoretically estimated synonymous transitions, ***Stv***_***o***_ is number of synonymous transversions observed, ***Stv***_***e***_ is the number of theoretically estimated synonymous transversions, ***Nti***_***o***_ is number of non-synonymous transitions observed, Ntie is the number of theoretically estimated non-synonymous transitions, *Ntv*_***o***_ is number of non-synonymous transversions observed, ***Ntv***_***e***_ is the number of theoretically estimated non-synonymous transversions. We calculated 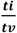 ratios for all the coding sequences using a program: a script written in Python-language is available at GitHub in the following link: (https://github.com/CBBILAB/CBBI.git).

Using our improved approach, the estimated *Sti, Stv, Nti*, and *Ntv* values for the 2516 coding sequences were calculated and analyzed. To further understand the better applicability of our approach, we found out 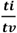 in all these 2516 coding sequences using maximum likelihood (ML) approach available in MEGA-X (Kumar *et al*. 2018) and compared it with the values estimated using the improved approach described in this study.

### Comparative representation of different 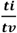 through the conventional and the improved estimators

A comparative analysis of 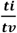 values calculated by using the conventional method and the improved method was performed to prove the better accuracy of our method proposed in this study. Also, we calculated the ratio between the *two-fold degenerate* and the *four-fold degenerate* codons (TFD: FFD) and the percentage of pretermination codons (PTC%) in each gene and performed statistical analysis between different parameters. The percentage (%) change in 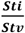 and 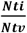 values obtained by conventional and improved methods calculated as follows.

a. 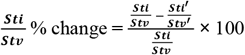
b. 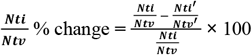

### Statistical analyses

OriginPro, Version 2022, OriginLab Corporation, Northampton, MA, USA was used to draw the Box-plot/Scatter plots as well as to perform the Mann-Whitney test (Mann and Whitney 1947) to find out the *p*-value. The correlation plot was drawn for R^2^ value and Pearson’s correlation coefficient (Pearson 1896) was also calculated between the 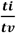 values obtained using the conventional and improved method.

## Results

### Impact of codon degeneracy and PTC on 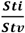 and 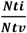 values in *E. coli*

We found out the total number of TFD, FFD, and PTC codons across 2516 coding sequences of *E. coli* and the percentage of TFD, FFD, and PTC was calculated (Supplementary Table 3). The percentage (%) compositional differences in the 2516 coding sequences represented as TFD%, FFD%, and PTC% were presented using box-plot (Supplementary Fig. 5). A variable range of TFD: FFD ratio was observed in the coding sequences of *E. coli*. The minimum to maximum TFD:FFD ratio was observed as 0.229 and 3.333 in *sugE* and *mntS*, respectively. A summary of the different parameters observed throughout the dataset was presented in Table 1. Among 2516 coding sequences, only 35 coding sequences possessed the TFD:FFD ratio of 1.000, which suggested that there exists a compositional asymmetry between TFD codon and FFD codon compositions across the coding sequences. We presented the TFD%, FFD% and PTC%, using a multi-paneled scatter plot to study the correlation between any two pairs (Fig. 2). As expected, the FFD and the TFD compositions exhibited a negative correlation (Pearson *r* value -0.56). The FFD and the PTC compositions exhibited a negative correlation (Pearson *r* value -0.65), while the TFD and the PTC compositions exhibited a positive correlation (Pearson *r* value 0.83) because the eighteen PTC codons include ten TFD codons and only one FFD codon. It also explained a positive correlation (Pearson r value 0.81) observed between TFD: FFD ratio and the PTC. The compositional variation (TFD, FFD and PTC) observed in coding sequences were considered to validate the accuracy of estimation of different 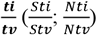 ratio using the conventional and the improved method.

**Fig. 2.**
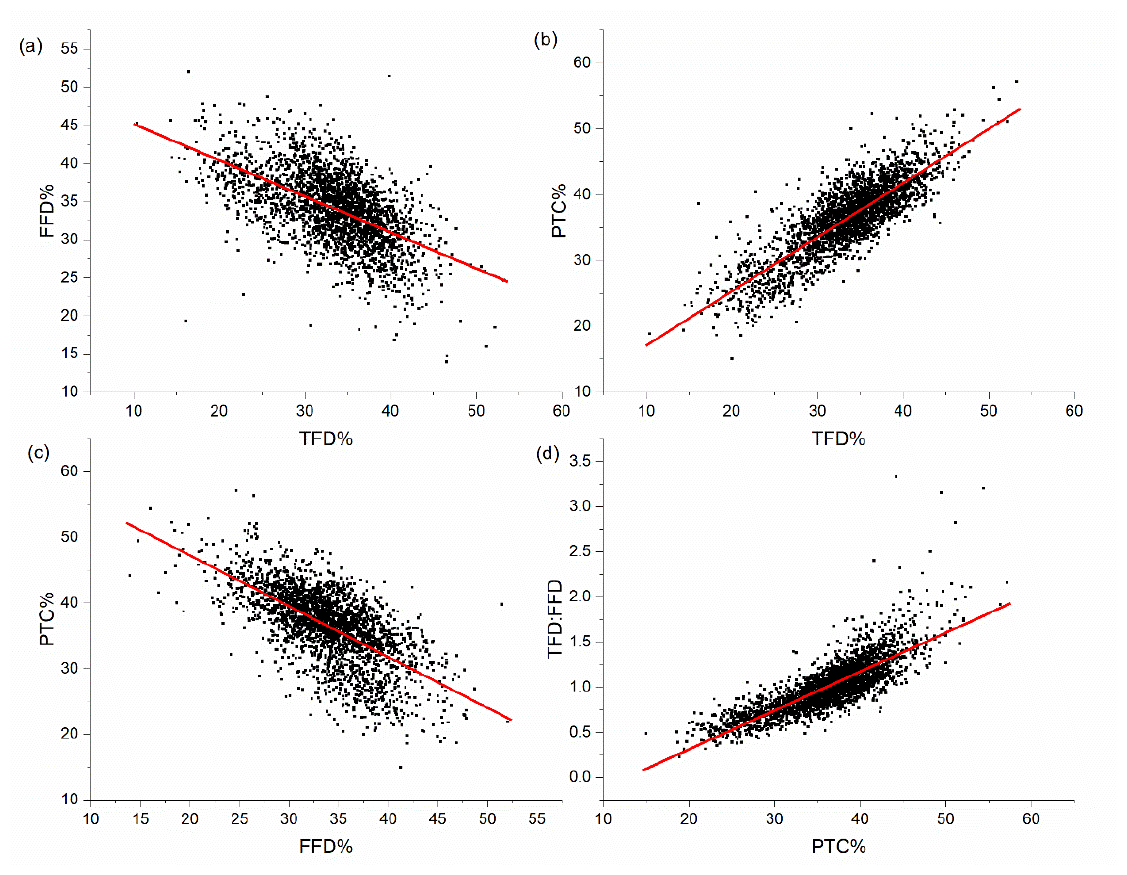
The multi-paneled scatter plots elucidate the comparison of codon composition between different parameters such as TFD% to FFD%, FFD% to PTC%, TFD% to PTC % and TFD: FFD to PTC%. In total 2516 *E. coli* genes were considered while calculating these values. The *x*-axes and the *y*-axes represent different parameters of codon compositions in the individual plots. The Pearson *r*(TFD%, FFD%) and Pearson *r*(FFD%, PTC%) were observed to be -0.56 and -0.65 respectively, indicating a negative moderate correlation. Whereas, the Pearson *r* (TFD%, PTC%) and Pearson *r*(TFD: FFD, PTC%) were observed to be 0.83 and 0.81 respectively, indicating strong correlation.

We estimated 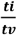 by accounting codon degeneracy as well as PTC in coding sequences and compared with the conventional method of 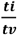 in coding sequences. Theoretically, a coding region entirely composed of FFD codons will result into a twice 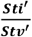 value than 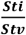 value, similarly a coding region composed of same proportion of FFD and TFD codons will result into equal values of 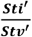 and 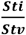. The increase in TFD codon proportion will have direct impact on the difference between 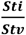 values and 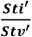 values. This pattern was observed in a hypothetical sequence considered as presented in the Supplementary Fig. 2.

The complete set of observed and estimated values of different variations in the 2516 coding sequences of *E. coli* were enlisted (Supplementary Table 3). These values were used to estimate the 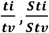 and 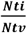 using the conventional method. The improved method was used to also estimated the 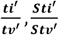 and 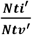. A comparative analysis was made between the values derived using the conventional method (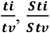and 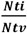) and the values derived using improved method (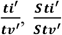 and 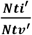) (Supplementary Table 3). A box-plot showing the comparison between the values of different 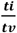 ratio obtained through the conventional and improved method is represented in Supplementary Fig. 7a. It was evident that the values estimated using the improved method were significantly different from that estimated using the conventional method (*p* value <0.01). The change in conventional and improved ratios between the synonymous and the non-synonymous 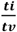 could be better visualized in Supplementary fig. 7b.

The synonymous variation in the coding sequences is likely to be different considering the differential composition of TFD and FFD codons, because when FFD codon proportion is higher in a gene, 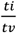 value is going to be lower. Similarly, when the FFD codon proportion is lower, 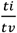 value is going to be higher. We analyzed the coding sequences exhibiting significant discrepancies between the two set of values 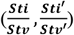. Some of the genes exhibiting higher value in the 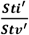 are DNA binding transcriptional regulation, ABC transporter, DNA polymerase III subunits, and Cytochrome Bo3 subunits. These coding sequences are comprised of higher FFD% than the TFD%. Some of the genes such as transcriptional regulator, toxin-antitoxin biofilm protein, and Iron-Sulphur cluster exhibited lower value in the 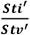 as these coding sequences are composed of higher TFD% than FFD %.

The Pearson correlation coefficient value between TFD: FFD and % change in 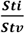 (conventional to improved) was obtained to be 0.891, signifying a strong correlation. Similarly, the Pearson correlation coefficient value between PTC% and % change in 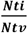 was obtained to be -0.488, a moderate negative correlation between the two variables. The results were found to be statistically significant at *p*<0.01, for 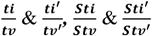 and 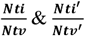. Further the correlation graph between % change in 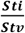 and TFD: FFD and % change in 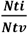 and PTC% also substantiate the role of codon composition and 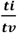 ratio (Fig. 3a & 3b). Further, we performed a simulation study to substantiate the impact of the compositional difference of degeneracy class in estimating 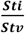 by using our improved method in coding sequences (Supplementary Table 8, Supplementary Fig. 3). The values obtained by the improved method were also different from the values obtained by the ML based method (Supplementary Table 9, Supplementary Fig. 4), as the degeneracy as well as PTC was not accounted in the ML based method.

**Fig. 3.**
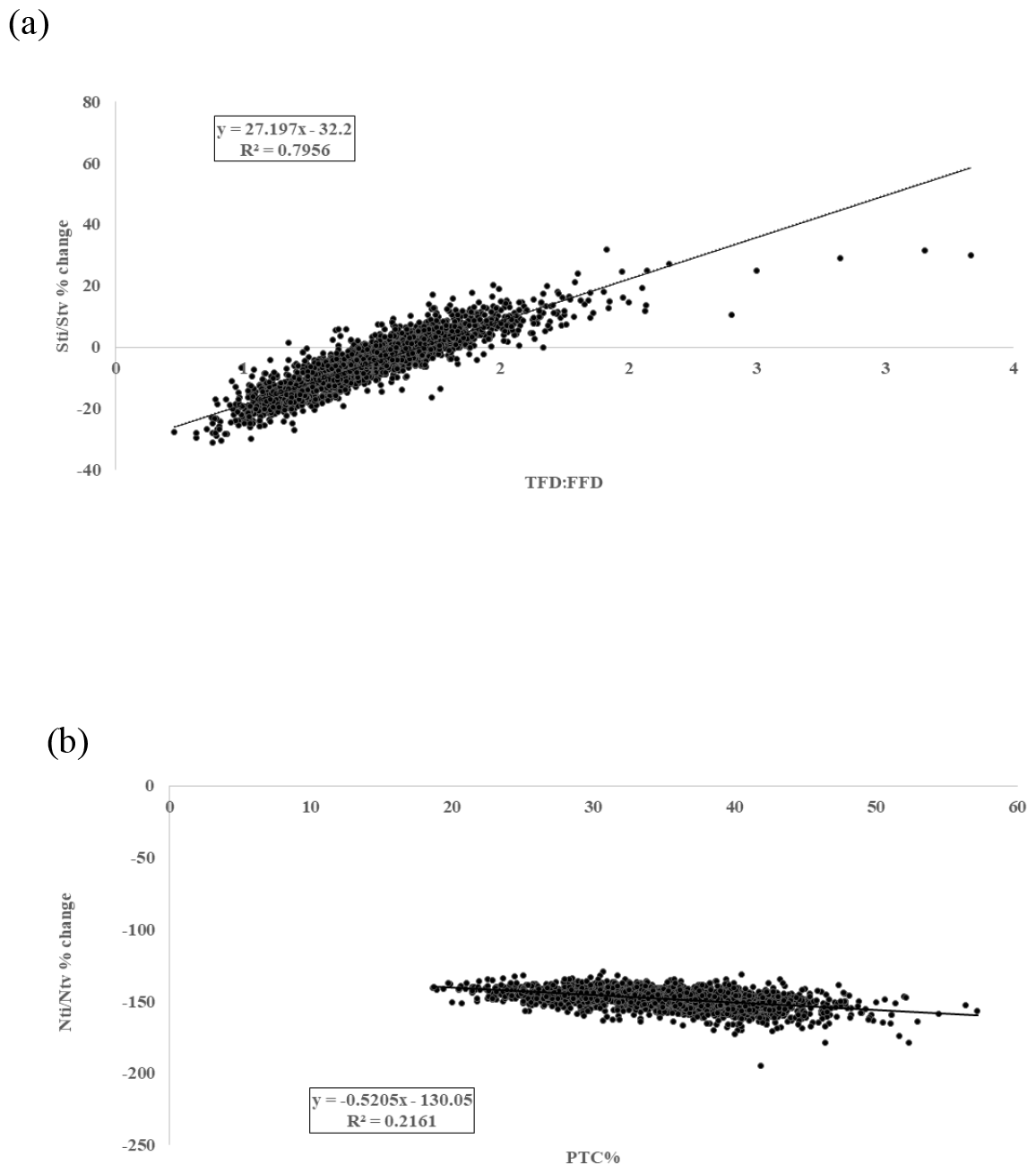
Regression plots between % change in 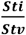 and TFD: FFD is presented in fig.3(a) and that between 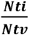 and PTC% is presented in fig.3(b) for the *E. coli* genes. Percentage change in 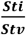 is showing a strong correlation with the TFD: FFD with a Pearson correlation coefficient value 0.89, whereas % change in 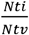 is showing a weak negative correlation with the PTC% with a Pearson correlation coefficient value -0.46.

**Fig. 4.**
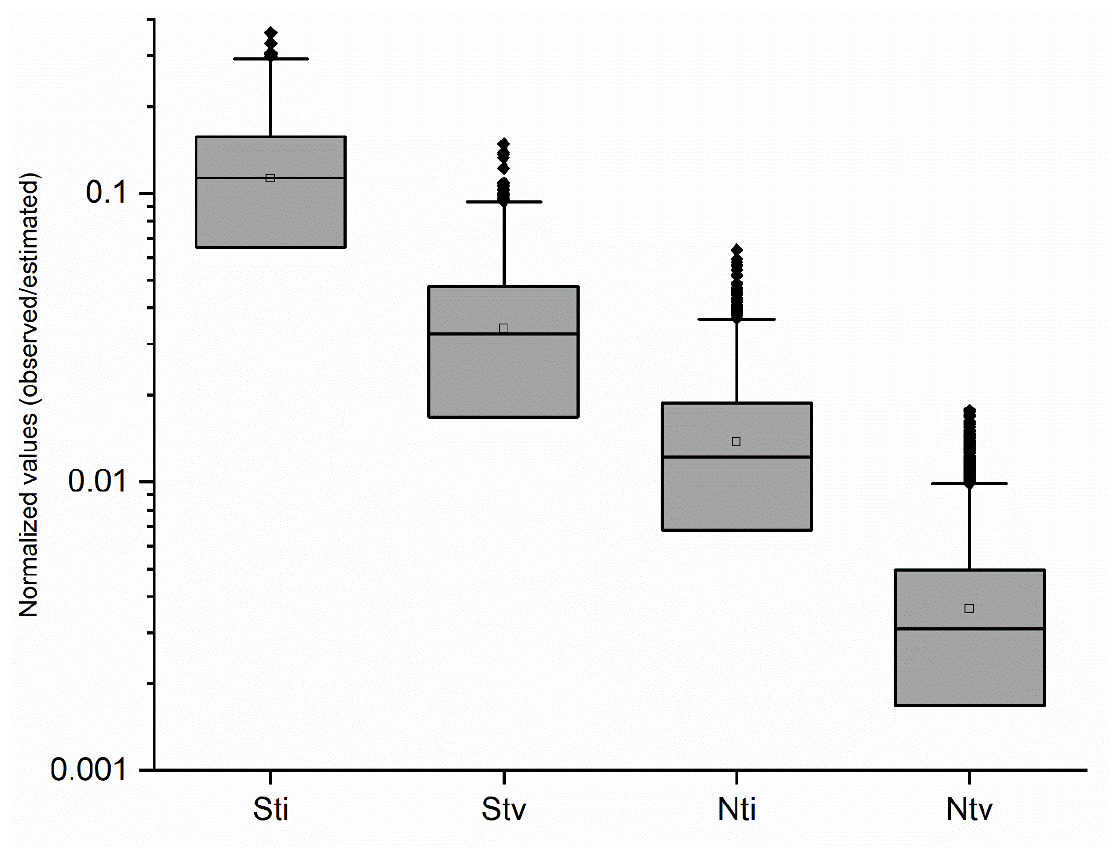
The figure illustrates distribution of *Sti, Stv, Nti*, and *Ntv* values using box-plot. The log scale has been used in the *y*-axis. The mean *Sti* and *Stv* were observed as 0.113 and 0.031 respectively, similarly the mean *Nti* and *Ntv* values were observed as 0.014 and 0.004. *Sti* was observed to be more than *Stv* and *Nti* was observed to be more than *Ntv*. The *Sti* and *Stv* were significantly different (*p*<0.01) from *Nti* and *Ntv* respectively.

**Fig. 5.**
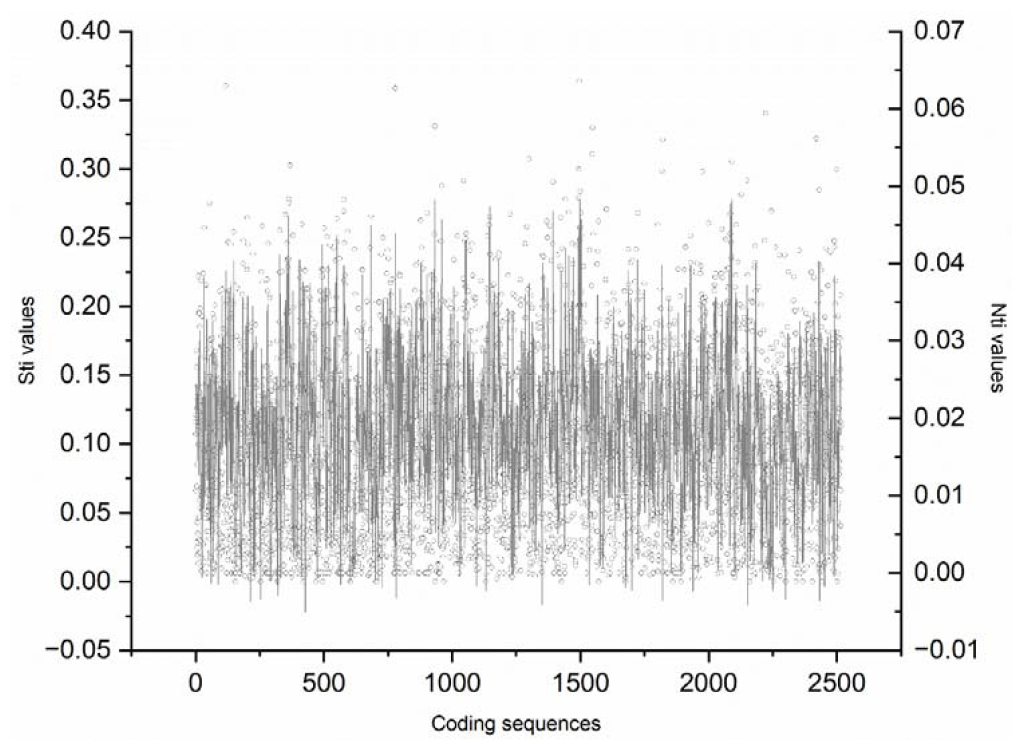
The figure illustrates the *Sti* and *Nti* values of coding sequences used in the study through a LOESS curve. The left *y*-axis shows the *Sti* values of coding sequences, and the right *y*-axis shows the *Nti* values of the coding sequences. The *x*-axis shows the numbers of coding sequences considered in this study. Each black bubble in the background shows the quantities of the ratio calculated for *Sti* and *Nti*. A Weak correlation (*Pearson r*=0.19) between the two variables can be visible distinctly throughout the SINE wave. It is evident that many coding sequences with similar Sti values have a wide range of Nti values.

### *Sti, Stv, Nti* and *Ntv* frequency in coding sequences

The frequency of *Sti, Stv, Nti* and *Ntv* were calculated and compared across the 2516 coding sequences. For Instance, the *Sti* frequency was calculated by considering the number of total *Sti* changes divisible by total possible *Sti* changes in a gene. Similarly, the remaining parameters (*Stv, Nti* and *Ntv*) were also calculated using the above logic, which follows our improved estimator i.e. observed number of changes out of total possible changes. The Supplementary Table 11 represents the overall *Sti, Stv, Nti* and *Ntv* values of the 2516 coding sequences considered for the study. The mean frequency values of *Sti, Stv, Nti*, and *Ntv* in the 2516 coding sequences were 0.113, 0.034, 0.014 and 0.004, respectively. (Supplementary Table 11, Fig. 4). This suggested the decreasing order of the different substitutions in genome such as *Sti* > *Stv* >*Nti* > *Ntv*. The values suggested that *Sti* was 3.3 times more frequent than *Stv, Nti* was 3.7 times more frequent than *Ntv, Sti* is 8.2 times more frequent than *Nti* and *Stv* was 9.3 times more frequent than *Ntv*. The relative purifying selection on *Nti* and *Ntv* in relation to their synonymous counter parts were similar. *Sti*, the most frequent substitution, was 28.25 times more frequent than *Ntv*, the least frequent substitution in this study. This we believe a distinct demonstration of frequencies of different substitutions such as *Sti, Stv, Nti*, and *Ntv* in *E. coli* coding regions.

Further we did a Pearson correlation study between different base substitution values that were obtained by using the conventional estimator as well as by the improved estimator (Table 2). The Pearson correlation value (*r*) between *Sti* and *Stv* was 0.920 in conventional method and 0.795 in improved method, respectively. Between *Nti* and *Ntv*, the Pearson correlation value was observed as 0.789 in conventional method and 0.612 in improved method, respectively. A lower correlation in the case of improved method was expected considering the normalization of coding region size in this method. Therefore, the correlation result using the values from the improved method was the true representation than the correlation values obtained in case of the conventional method where size of the coding region was not normalized. This was more evident in the correlation between *Sti* and *Nti*: the Pearson correlation value (*r*) between *Sti* and *Nti* was 0.527 in conventional method whereas the same was 0.192 in case of the improved method. The low correlation is expected here due to the independent nature of the purifying selection on non-synonymous variation. Therefore, our improved method is more appropriate in estimating the distinctive selection patterns on coding regions by doing comparison between *Sti* and *Nti* (Table 2).

As part of our comparative study, we investigated the coding sequences of similar *Sti* and different *Nti* values (Fig. 5). In a set of 20 coding sequences with similar *Sti* values such as 0.131, the *Nti* values were ranging from 0.000 to 0.030. This wide range of *Nti* values, with a maximum 30-fold variation, highlighted the presence of a variable *Nti* pattern within the respective coding sequences. Similar observation could be found in other *Sti* values. We compared *Sti* and *Nti* values in relation to gene expression represented with codon adaptation index (CAI) values (Sharp and Li 1987; Sen et al. 2019). We studied coding sequences with size more than 500 bp. The top 100 coding sequences with maximum *Sti* values (mean 0.255) were compared with bottom 100 coding sequences with minimum *Sti* values (mean 0.006). The mean CAI values for the top 100 coding sequences were 0.565 whereas the mean CAI values for the bottom 100 coding sequences was 0.509. The two mean CAI values were found to be close. This indicates the *Sti* values might not be influenced significantly due to gene expression. Similarly, the top 100 coding sequences with maximum *Nti* values (mean 0.036) were compared with bottom 100 coding sequences with minimum *Nti* values (mean 0.001). The mean CAI values for the top 100 coding sequences were 0.491 whereas the mean CAI values for the bottom 100 coding sequences was 0.638. The two mean CAI values of the coding sequences were found to be significantly different (*p*<0.01). The selection due to gene expression is stronger on *Nti* than on *Sti* in *E. coli*.

## Discussion

Codon degeneracy influences the *ti* to *tv* ratio in coding sequences because a TFD codon undergoes synonymous variation only by transition while a FFD codon undergoes synonymous variation by transition as well as transversion (Aziz *et al*. 2022). In addition, there are only 18 pretermination codons in the genetic code table where nonsense substitution is rare to observe. Coding sequences in a genome vary with composition of codons belonging to different degeneracy as well as PTC. Therefore, it is important to compare coding sequences regarding their 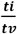 ratio by accounting degeneracy and PTC. Accordingly, in this study we have developed a methodology to estimate 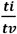 ratio by accounting to the above features in coding sequences. In total 2516 coding sequences of *E. coli* have been analyzed. The impact of codon degeneracy and PTC is observed distinctly on the ti/tv values in coding sequences. Interestingly, the coding sequences are observed to be different regarding their 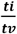 ratio even after accounting to the composition of degenerate codons and PTC. In fact, the 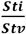 values are observed to be variable among the coding sequences, which is intriguing. We did not observe any significant correlation of 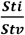 values with the gene expression represented as CAI values. The role of context dependent mutation (Sung *et al*. 2015; Zhu *et al*. 2017; Aikens *et al*. 2019; Ling *et al*. 2020), which has been described recently, might be a contributing factor that needs to be investigated in the future.

Our investigation into *ti* and *tv* revealed that *Sti* is around three times more frequent than *Stv* while it is eight times more frequent than *Nti*. The difference between *Sti* and *Stv* may be attributed due to the structural changes in DNA happening due to mispairing between either purine: purine or pyrimidine: pyrimidine but the difference between *Sti* and *Nti* may be attributed to the purifying selection acting on the coding sequences because of the non-synonymous changes. The magnitude of the purifying selection acting on non-synonymous substitution in *E. coli* genome is vividly observed. The strength of purifying selection on *Nti* and *Ntv* seems comparable considering the closeness of the 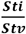 and the 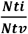 values. In a recent study by Zou and Zhang (Zou and Zhang 2021) suggests that the purifying selection is more on *Ntv* than on *Nti* which is unlike the observation reported in our study. Our different observation from that of Zou and Zhang might be attributed to the intra-species study limited to *E. coli* whereas Zou and Zhang worked on an inter-species approach covering 90 clades representing all domains of life. Further, in our estimation of *Nti* and *Ntv* we have accounted codon degeneracy and PTC.

The correlation values between different substitutions such as *Sti* and *Stv*, as well as between *Sti* and *Nti* were higher in case of the conventional method than that in the case of the improved method. This is an important observation in our study and distinctly demonstrate the superiority of the improved method over the conventional method, which is explained as follows. In case there is no selection on *Sti*, the number of *Sti* is likely to be directly proportional with the number of synonymous transition sites available in a coding sequence. So, *Sti* is likely to correlate strongly with gene size. In case of *Nti*, purifying selection is strong, for which the number of *Nti* observed in a coding sequence is significantly lower than the number of *Sti* observed in the sequence though the number of theoretically possible *Nti* sites is always higher than the number of theoretically possible *Sti* sites. The number of observed *Nti* is also likely to be proportional with the possible *Nti* sites in a coding sequence. So, the number of *Nti* is likely to be proportional with gene size. Therefore, we can anticipate a strong positive corelation between *Sti* and *Nti* in the 2516 coding sequences. This is indeed what we have observed in our analysis here. In the improved method to estimate *Sti* and *Nti* described in this study has normalized the number of possible sites for *Sti* as well as *Nti* (or gene size). The improved *Sti* approach provides us an idea regarding the frequency of the substitution per *Sti* site whereas the improved *Nti* approach provides us an idea regarding the frequency of the substitution per *Nti* site. As size has been normalized, corelation between improved *Sti* and improved *Nti* is likely to be lower. Further, the value of improved *Sti*/improved *Nti* is more suitable to understand the purifying selection on a gene than *Sti*/*Nti*. A similar explanation may be also given for the *Sti* and *Stv* correlation. As gene size increases, *Sti* and *Stv* also increases, for which strong positive observed between the between the two values. In case of improved *Sti* and improved *Stv* cases the possible sites have been normalized respectively. Susceptibility of different sites for *ti* and *tv* are not same as evident by C→T *ti* and G→T *tv* are more frequent than the other *ti* and *tv*, respectively (Sen *et al*. 2022; Beura *et al*. 2023). Therefore, the correlation value between improved *Sti* and improved *Stv* is likely to be weaker than the same between *Sti* and *Stv* values.

As transitions are more frequent events than transversion, transition can be considered alone to understand the selection on coding sequences. Considering *Sti* representing the variation frequency in a coding sequence, the *Nti* representing the strength of purifying selection acting on the coding sequence, a comparison between *Sti* and *Nti* in a coding sequence might represent the strength of purifying selection acting on it. In several instances, we have observed that coding sequences with equal *Sti* variation frequency are highly different regarding their *Nti* variation frequency, which indicates disproportionate purifying selection on coding sequences. Additionally, when we compared the mean CAI values of the coding sequences with higher *Sti* values against the mean CAI values of the coding sequences with lower *Sti* values, we did not find any significant difference. On the contrary, when we conducted the same analysis for the *Nti* values, we observed a significant distinction between the CAI values of coding sequences with higher *Nti* values and those with lower *Nti* values. This observation indicates that highly expressed genes tend to have lower *Nti* values. Consequently, it suggests that purifying selection due to gene expression is more pronounced for *Nti* values than for *Sti* values. Since our study on selection relies on intra-species analysis, we anticipate that future research will unveil the underlying mechanisms of selection governing variation frequency in inter-species studies as well. A probable discrepancy between the process of selection between intra-species and inter-species can be drawn. This is a simple demonstration of observing purifying selection acting on coding sequences. Though the difference between *ti* and *tv* have been known for a long time, we could use it to study selection on coding sequences because of the improved estimator.

Various methods have been implemented in the estimation of the *ti* to *tv* ratio by molecular evolutionary biologists to understand the mutation as well as selection bias. The phylogeny-based method and Bayesian methods have been implemented by researchers in the inter-species study of 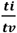 (Purvis and Bromham 1997; Huelsenbeck *et al*. 2001). Usually the *dN*/*dS* approach has been used to visualize the selection in coding sequences (Aziz *et al*. 2022) that considers both *ti* and *tv*. However, our method implemented in this study is simple, succinct, easy to follow for researchers in the field with limitations in mathematics and provides an insight to understand the evolution of protein coding sequences. We also did a comparison with the ML method. It has been said that the likelihood methods are computer intensive and difficult to apply to large datasets and can fall into local traps (Golding and Felsenstein 1990). Furthermore, the extrapolation of our method in various inter-species studies, as well as phylogenetic tools can enhance the horizon of our understanding regarding mutational bias in coding sequences.

## Data and software availability

A script written in Python-language is available at GitHub in the following link: (https://github.com/CBBILAB/CBBI.git). Further queries regarding the software may contact SSS. The authors confirm that the data supporting the findings of this study are available within the article and Supporting information.

### Acknowledgments

We are thankful to the anonymous reviewers and the editors for their critical suggestions to improve the quality of the manuscript. PKB is grateful to Tezpur University and DBT project grant (BT/PR40231/BTIS/137/63/2023) for the fellowship. RA is thankful for the JRF fellowship from the DBT grant (BT/511/NE/TBP/2013) and (BT/403/NE/U-Excel/2013). PS is grateful to UGC, GoI New Delhi, for the JRF. SD is thankful to DBT for MSc. Scholarship. SSS and SKR are thankful to DBT, GoI for the twinning grant (BT/PR16361/NER/95/192/2015 date 18-10-2016) as well as the DBT project grant (BT/PR40231/BTIS/137/63/2023) to them. NDN, SSS & SKR thankfully acknowledge the DBT funded Bioinformatics and Computational Biology Centre at Tezpur University (BT/PR40253/BTIS/137/52/2022). We are also thankful to Prof. Kini R Manjunatha, National University of Singapore for his kind suggestions during his visit to Tezpur University.

